# Integrative systems approach reveals dynamics of microbiome-metal-ion axis in mesocosms representing tropical urban freshwater canal ecosystem

**DOI:** 10.1101/2020.09.21.306803

**Authors:** Gourvendu Saxena, Eric Dubois Hill, Ezequiel M. Marzinelli, Shivshankar Umashankar, Toh Jun Wei, Wei Woo Yissue, Peter D. Steinberg, Verónica B. Rajal, Staffan Kjelleberg, Rohan B. H. Williams, Stefan Wuertz, Sanjay Swarup

## Abstract

Freshwater ecosystems of <u>tr</u>opical <u>u</u>rban <u>c</u>anals <u>s</u>ystems (TrUCS), are highly dynamic and experience constant pressures from interspersed effects of land-use and rain. The dynamic nature of TrUCS ecosystems presents a unique opportunity to unravel the signature interactions between the macro-organisms (top-down), <u>sed</u>imentary <u>mi</u>crobial <u>c</u>ommunities (SedMICs), their functioning and the geochemical environment (bottom-up). A systems level understanding of the molecular and mechanistic basis of the highly dynamic behaviour that leads to specific ecosystem outcomes, is currently lacking. Therefore, a research framework to identify the direct link between top-down and bottom-up ecological effects on SedMICs in a highly dynamic urban canal sedimentary system is needed. Here, we present a framework of integrated multi-dimensional data across system-level biotic and abiotic ecological descriptors, such as environmental variables and active SedMICs. We followed the ecosystem shifts after a natural disturbance (rain) in two different anthropogenic disturbance (land-use) regimes. Shifts in profiles of metabolically active community were conserved across different land-use types, indicating resilience to perturbation is an intrinsic property of the TrUCs ecosystem. Three distinct phases, which were dominated sequentially by autotrophy, anoxic-heterotrophy and oxic-heterotrophy, were identified within these shifts. The first two phases were influenced by the bottom-up effects of specific metal-ion combinations of nitrates and sulfates with magnesium, aluminum and iron, and the third phase was triggered by top-down influences of bioturbation. This generalized systems-level approach, which provides an ecosystem-centric understanding of TrUCS and integrates them in sustainable management practices, can also be extended to other freshwater ecosystems.

## Introduction

<u>Tr</u>opical <u>u</u>rban <u>c</u>anal <u>s</u>ystems (TrUCS) are unique and dynamic ecosystems, driven by both natural and anthropogenic controlling factors that can influence both top-down effects, such macro-ecological bioturbation and predation, and bottom-up effects, such as nutrients transformation on the microbial ecosystem. The complex <u>sed</u>imentary <u>mi</u>crobial <u>c</u>ommunities (SedMIC) in TrUCS ecosystems with inherent self-cleaning potential are shaped by these factors these factors ^1, 2^. Yet, the interactions between microbial communities and the ecosystem controlling factors at a systems level are poorly understood. Consequently, the direct link of combined top-down and bottom-up effects on SedMICs in a highly dynamic urban canal sedimentary system is not known. Therefore, attempts to manage the urban canals in an environmentally sustainable way do not incorporate an understanding of how microbial processes interact with the controlling factors of urban environments.

Environment controlling factors, such as land use and rain represent chronic anthropogenic and intermittent natural disturbances, respectively ^3–5^. The constant release of land-use-specific stressors through basal flow of water exerts a chronic anthropogenic (press) disturbance in the sedimentary ecosystems of TrUCS ^1^ from different land use types. For example, while organics enter the canal ecosystem mainly via increased sediment flow, the water-soluble components, such as nutrients, enter the canals through the water phase ^1^. A diverse pool of metals entering the TruCS has the potential to influence the functioning of microbial communities ^1^, specifically, in assisting the energy harvest processes. Nutrient transformations, on the other hand, can have a profound effect on primary productivity and heterotrophic processes in TrUCS. The involvement of metals such as aluminum in shaping the functions of SedMICs ^1, 6^ indicates the presence of novel interactions and associations of environmental constituents with energy and nutritional pathways. The outcome of these interactions might have a direct effect on surface water quality.

Rain represents a relatively low-severity, high-frequency intermittent natural disturbance, particularly in tropical regions. It can bring rapid changes in the levels of physicochemical stressors, such as frequency of dry-wet cycles and levels of metals, organics, antibiotics and emerging pollutants ^5^, which could have a major influence on the resident SedMICs. Rainfall can be as high as 10 mm/day during the wet season and 4 mm/day even during the dry season ^7^. Tropical and subtropical rain events and dry periods can cause intermittent deep and shallow water in storm-water canals. The duration of resulting shallow water can vary from a few hours to weeks ^8^, with typically a one-month dry period between rain events. The hydrodynamics of concrete-paved waterways, constructed to rapidly convey water from the catchment ^9, 10^, adds to the fluctuating dry-wet conditions. Rain perturbations also directly influence instantaneous sediment concentration, which typically reach the levels of 2100 mg/l ^11^ in TrUCS.

The combined effects of chronic anthropogenic and intermittent natural disturbances have been understudied ^12^, despite their possible role in shaping the sedimentary ecosystems of TrUCS. The effects of land use and rain are not independent of each other. Hence, multi-scale integration of anthropogenic (for example, land use) and natural (for example, rain) controlling factors with ecosystem processes is required for a holistic understanding of TrUCS ecosystems, which to the best of our knowledge is still lacking. An understanding of ecosystem interactions on the molecular level (both at the level of ecosystem quality parameters and SedMICs gene-level communal response), leading to specific ecosystem outcomes is also lacking. Temporal patterns of shifts in the community composition and ecosystem functioning, triggered by intermittent or pulse disturbances, such as rain or fire, have generally been reported in macro-scale ecosystems, for example forests ^13^ and natural river systems ^14^. The ecological theories of successions derived from studying such macro-ecosystems indicate the dominant role of primary productivity, driven mainly by phototrophs ^13^ in the initial phases and then by heterotrophic processes ^13^. Such shifts occur over timescales of months and decades in ecosystems such as forests and natural rivers ^13^. In microbial ecosystems, however, the generation time of microorganisms is significantly shorter, compared to that of the macro-organisms of larger ecosystems. Therefore, for microbial systems the longer temporal scales in such studies usually do not resolve the highly dynamic nature of compositional and functional shifts immediately following rain events and remain under-explained. Microbial ecosystems, such as SedMICs, are easily tractable ^15, 16^ and highly dynamic; hence they present a unique opportunity to test ecological theories of disturbances and temporal shifts on relatively short timescales.

To identity the combined role of chronic anthropogenic and intermittent natural disturbances on TrUCs sedimentary ecosystems, we used a mesocosm flume system that mimics a controlled linear channel flow tropical urban canal ecosystem. We thus unravelled a holistic understanding of the role of natural and anthropogenic controlling factors on ecosystem process dynamics. The influences of these controlling factors are integrated with the response of SedMICs on multiple scales, ranging from molecular pathways (bottom-up interactions) to the role of macro-organisms (top-down interactions) in the ecosystem. We first examined the temporal shifts in response to changes in major ecological processes, such as net primary productivity and heterotrophy, with respect to carbon and nutrient metabolism. We then identified the metal and nutrient interactions during biogeochemical temporal shifts and their associations with microbial communities and gene expression. Finally, we identified the members of microbial functional guilds involved in these transformations, thus outlining the stressors-species-genes relationships. By following the ecological events after rain in two different land-use types, we have described the potential mechanistic basis of interactions between natural intermittent and anthropological chronic disturbance with SedMICs of a TrUCS ecosystem. These outcomes can be further tested with individual microbial species in multiple controlled experiments to validate the findings. The results presented here will bridge gaps in our understanding on the mechanistic basis of sedimentary microbial ecosystem shifts in TrUCS.

## Results

A mesocosm system with linear-flow and partial replacement was designed to study the interactions of sedimentary microbial communities (SedMIC) with the associated ecosystem descriptors (Figure 1A). The study was conducted using the sediments from two land-use types, residential and industrial (Figure 1B), along with the water taken from the respective sites over a period of 30 days (Figure 1C). Sediment color (Figure 2A) changed from brown to black from day 0 to day 8, indicating a shift from an oxic to anoxic phase. It resumed a brown color over the 30-day experiment, indicating the reversal of sedimentary oxidative conditions. Macro-fauna activity (Figure 2B) was observed in the later phases, leading to prominent bioturbation. The macro-fauna was active in the upper and middle layer of sediments, which contributed to an overall oxidative environment through oxygen transfer.

**Figure 1:**
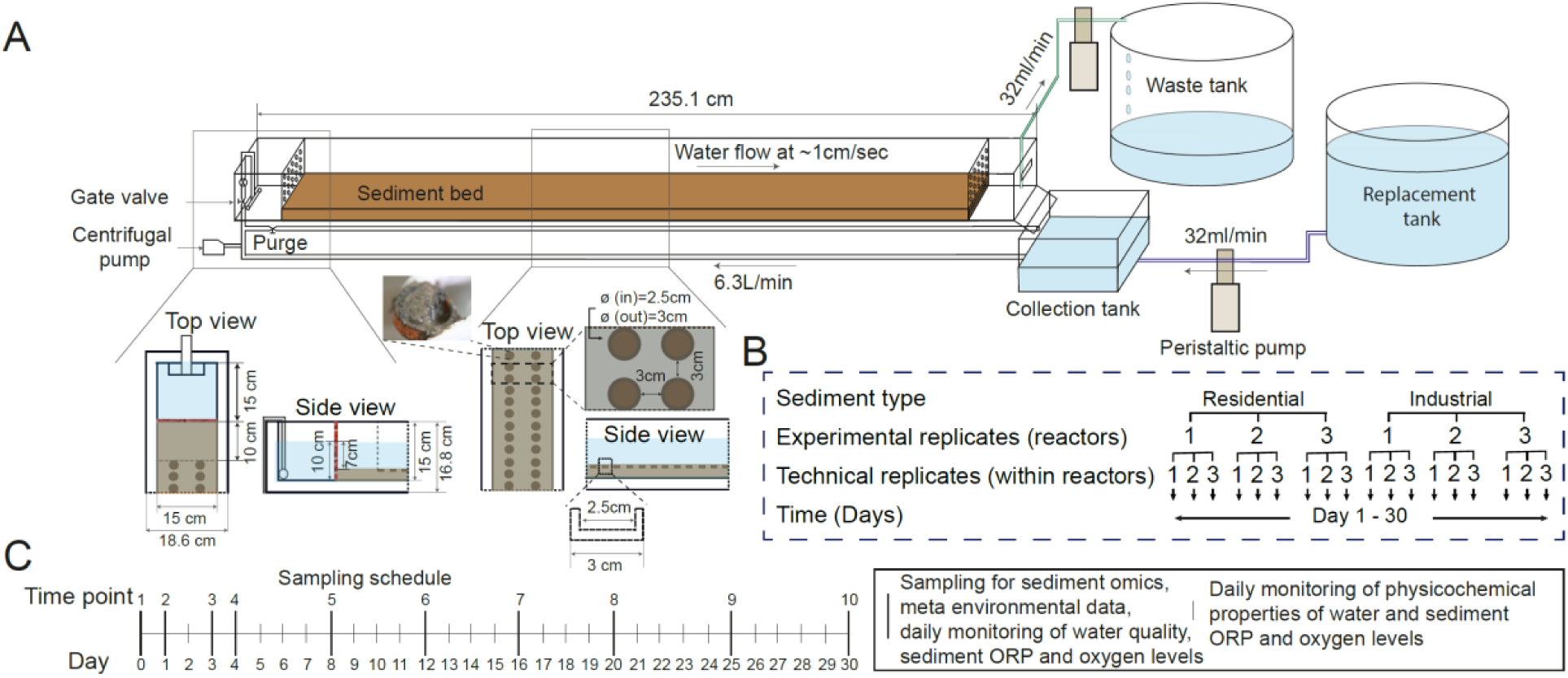
Schematic of the flume set-up and experimental design. (A) The flume is shown with recirculation and partial water replacement schemes set-up to mimic linear flow. The magnified sections provide the details of top and side view from the upstream and mid-stream sections of the flume system. (B) Experiment design with sediment-type (land-use) and time as fixed factors and experimental replicates as random factor. (C) The distribution of sampling time-points for data collection.

**Figure 2:**
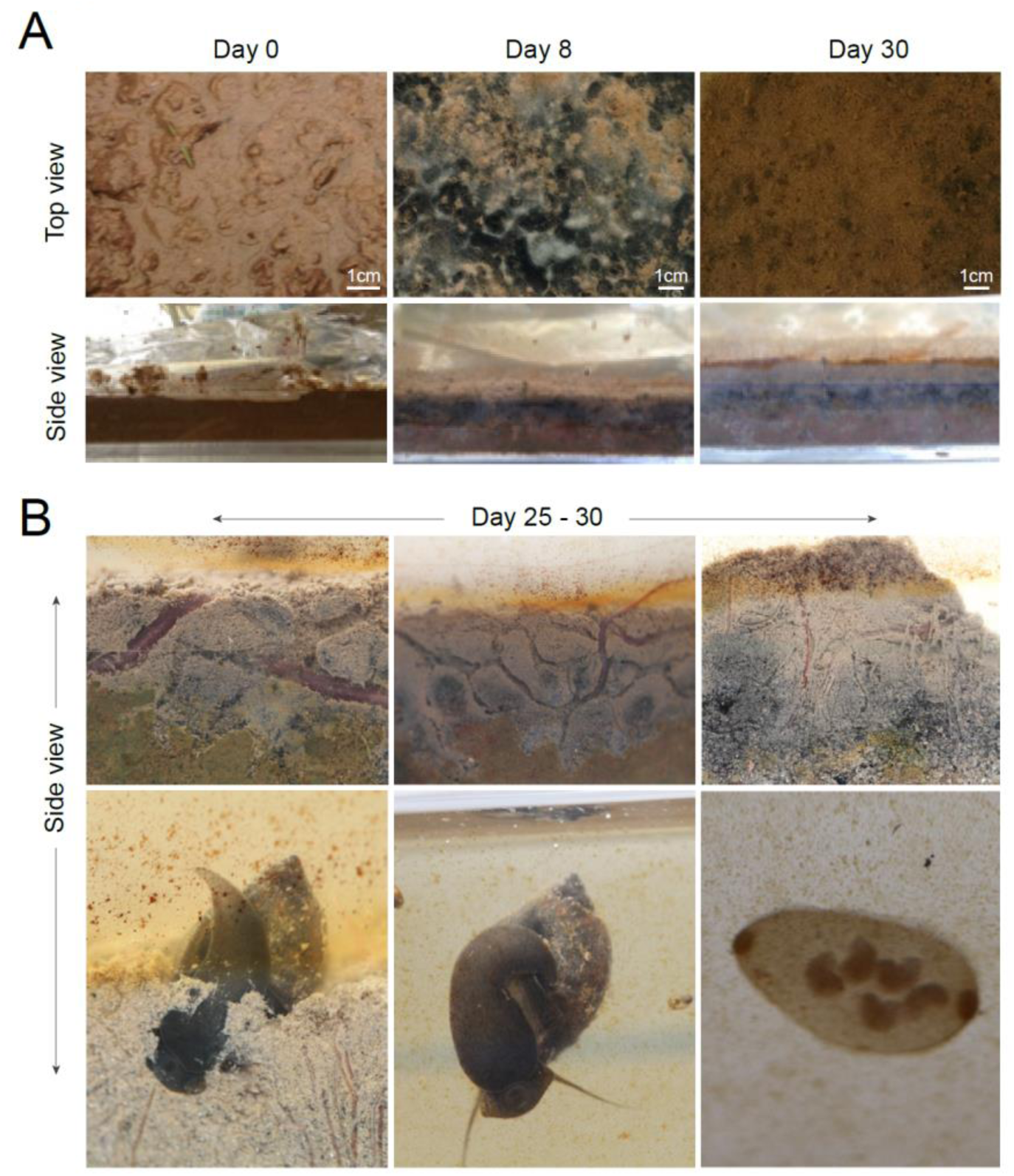
Sediment texture and macro-ecology in the mesocosm (A) Sediment texture is shown from early, mid and late time-points in the top panel from left to right. Texture shows dynamic shifts turning from brown to black and then to stable brown again. The bottom panel shows the depth profile of sediment texture. The black anoxic layer starts building during mid-phase of experiment, which is then covered by the brown layer of sediments in the late-phase of experiment. (B) Trophic levels preserved in the mesocosm flumes. The flumes were able to keep the dynamic macro-ecology alive during the 30-day experiment. The small worms help in oxygen transfer to the sediments by their sinusoidal motion. The big worms also contributed by burrowing and emerging from the sediments. Snails were spotted frequently grazing on the top of sediments.

### Ecological phasing in TrUCs ecosystem

Using unconstrained clustering analysis, the ecosystem descriptors indicated the presence of three distinct temporal phases (Figure 3A). Hierarchical k-means clustering (hkmeans) delineated the three phases into five different trends of ecosystem descriptors (Figure 3B and 3C). The observed regime shift in microbiota and functional guilds reflected the shifts in ecosystem descriptors. These three phases were classified as acclimation autotrophy phase, anoxic heterotrophy phase and oxic heterotrophy phase. In the later phases, bioturbation (mixing of sediment) by macro-fauna contributed to in creating an oxidative environment. The three phases based on shifts in ecosystem descriptors, SedMICs community and active functional guilds are explained in detail in the following sections.

**Figure 3:**
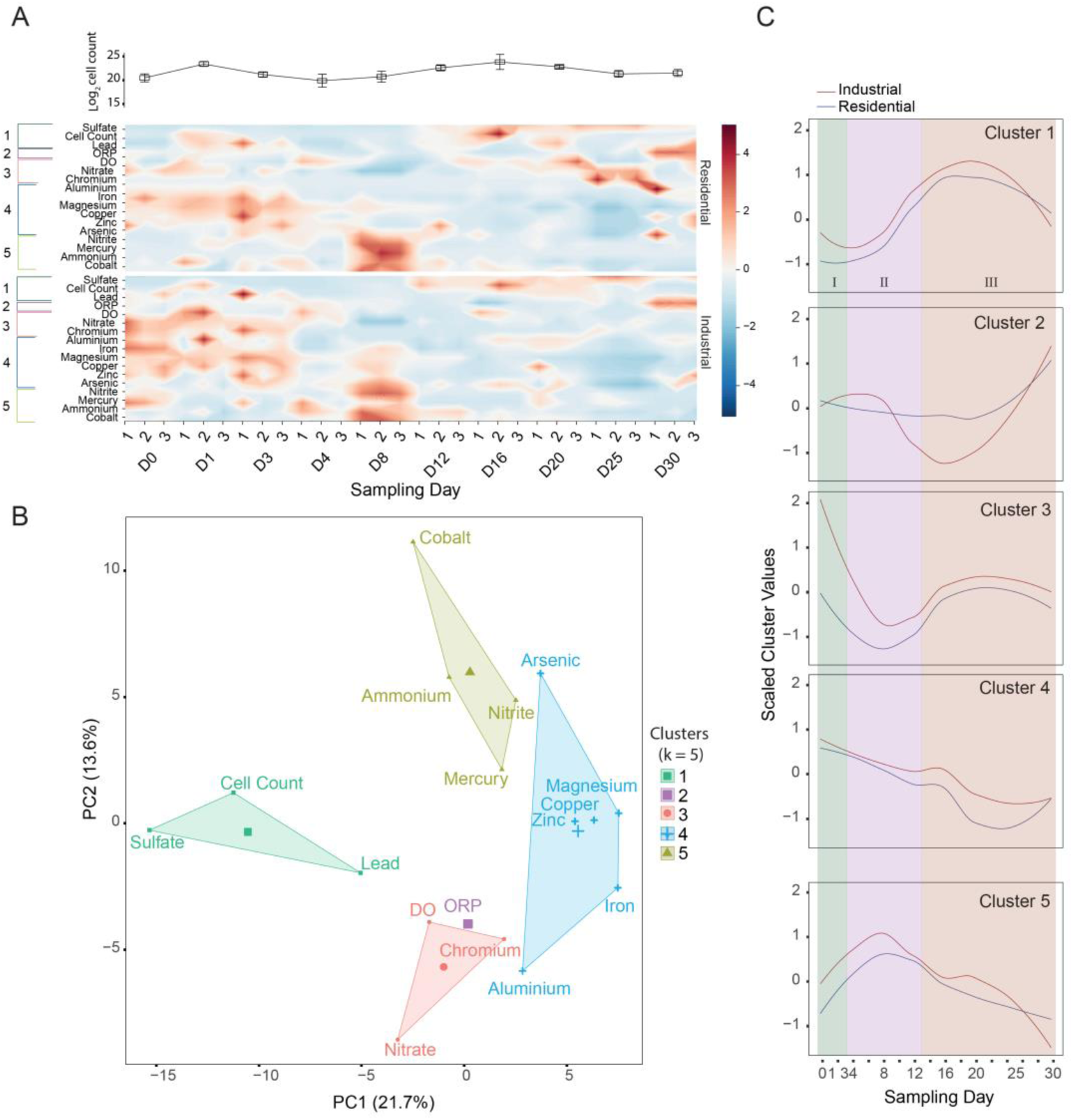
Relationships between ecosystem descriptors from sediments. (A) The ecosystem descriptors are plotted as continuous surface plot to highlight the temporal variations in their levels from day zero (D0) to day 30 (D30). The levels of ecosystem descriptors are mean-centered and scaled between −5 and +5. (B) The data was then clustered using hierarchical k-means clustering groups with similar trends. (C) The scaled average cluster values of ecosystem descriptors are plotted for residential and industrial land-use.

#### Shifts in the Ecosystem Descriptors of TrUCs ecosystem

The hierarchical k-means clustering (hkmeans) delineated three phases (Figure 3A) and displayed five characteristic trends of ecosystem descriptors (Figure 3B and 3C). In the acclimation autotrophy phase, which lasted from day 0 to 3, high levels of metals and oxidized forms of nutrients (Figure 3A) reflected the disturbance of sediments due to the shear-force of rain. As the sediments acclimated to the new conditions after rain, dissolved oxygen (DO) and oxidation-reduction potential (ORP) decreased, reflecting an anoxic/reducing environment of the anoxic heterotrophy phase (days 4 to 12), which resulted in increased levels of reduced nutrients and decreased levels of metals. In the oxic heterotrophy phase from day 12 onwards, the top-down forces of bioturbation led to increased oxygen transfer, as indicated by a high DO and ORP. Thus an increase in the levels of oxidized nutrients was observed (Figure 3A). The ecosystem descriptors from concordant water and sediment samples showed similar temporal trends in both land-use types (Figure S1). This observation provided an early estimate of similar pressures from the environment in different TrUCS.

#### Three phases of compositional and functional regime shifts identified in TrUCs ecosystem

The composition and active functional guilds of the microbial communities closely followed the patterns of temporal shifts, which were delineated into the three phases, as observed in ecosystem descriptors and sediment phenotypic changes. Despite the differences at each temporal stage, the trends of composition and functional shifts in SedMICs (Table 1A and B) were conserved for residential and industrial land-use types (Figure 4Ai and Aii). Community dynamics showed uniform dispersion over time with a clear pattern of temporal shifts from day 0 to day 30, along the primary axis (Figure 4Ai).

The top 15 phyla contributed to 20% of the variation and were associated with the temporal shifts in the SedMICs of the two land-use types (Figure 4B). Differential analysis of the taxonomic abundance data over time corroborated the association of these 15 phyla with the temporal shifts.

**Table 1:**
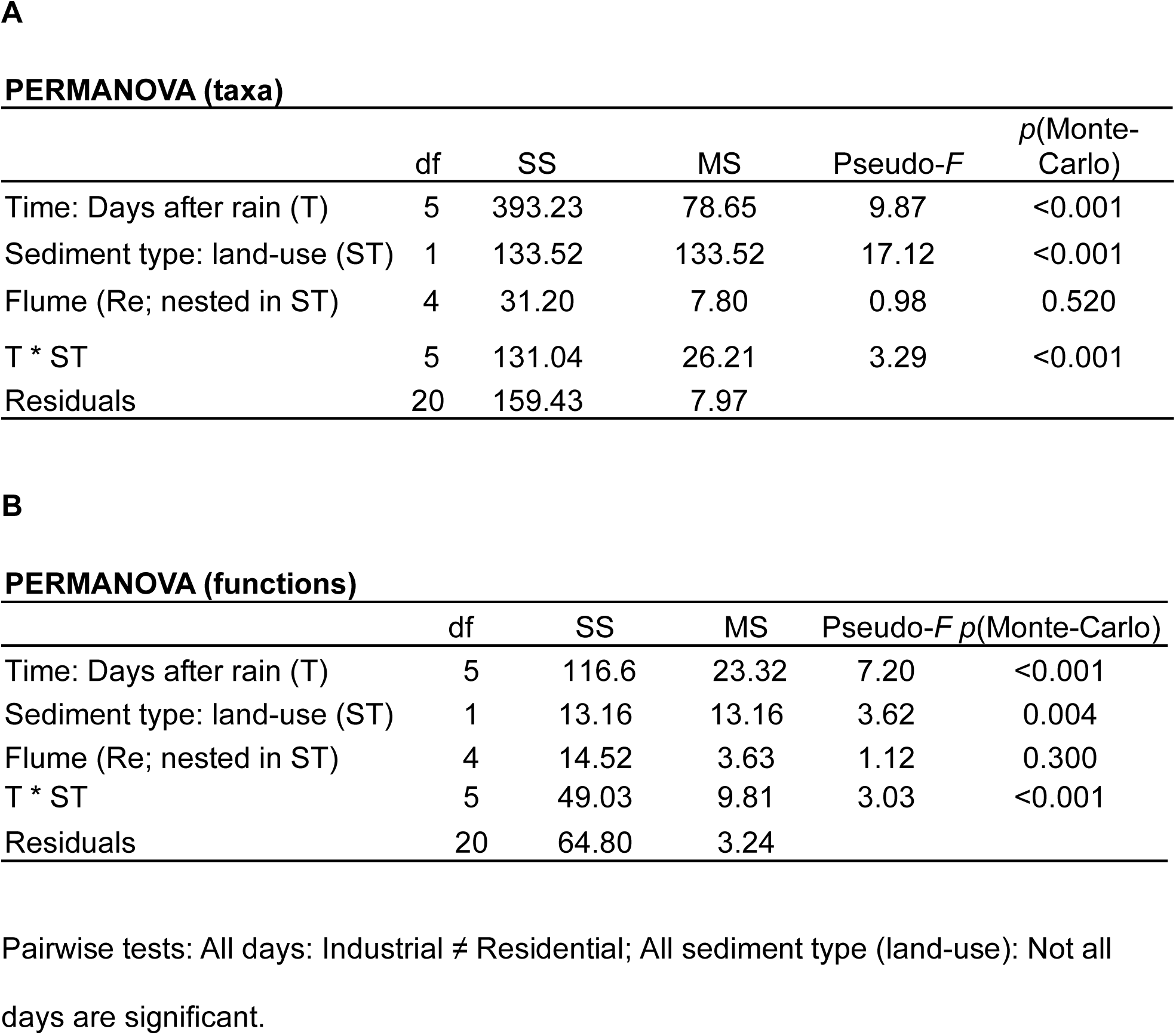
PERMANOVA results for active microbial communities’ profiles, using Bray-Curtis distance similarity on regularized logarithm transformed (rlog) abundance data from (A) active microbial taxa groups and (B) microbial functions. Time and sediment type (land-use) were the fixed and orthogonal factors. Flume number was random factor, nested in sediment type (land-use). ST: sediment type (land-use); T: Time (Days after rain); Re: Flume; df: degree of freedom; SS: sum of squares; MS: mean sum of squares; Pseudo-*F* = *F* value by permutation

**Figure 4:**
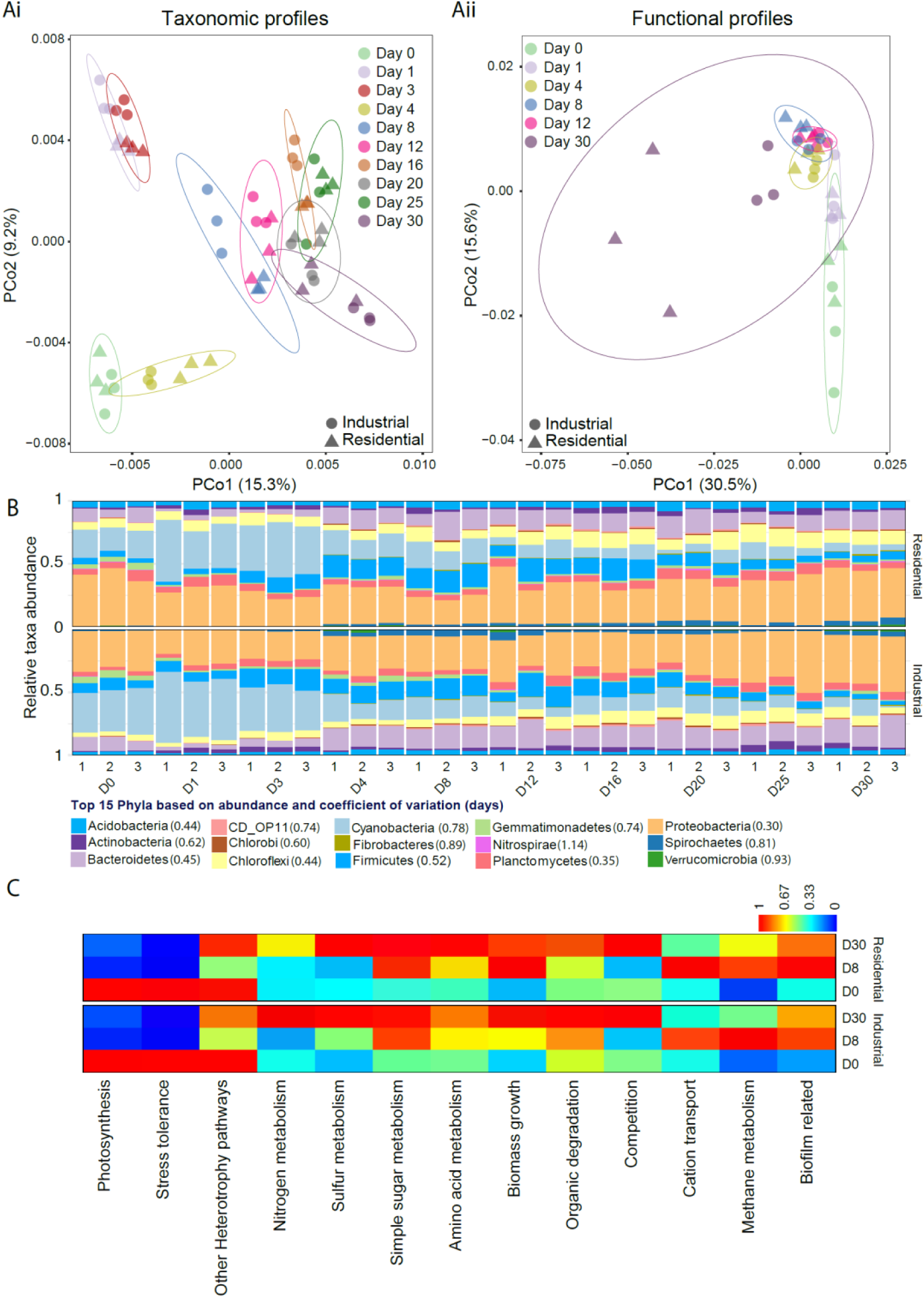
Temporal shifts in active microbial communities’ profiles. PCoA plot, showing the relative shifts in (Ai) active microbial taxa and (Aii) functional profiles of sedimentary microbial communities in urban waterways. Bray-Curtis distance was calculated to create similarity matrix for respective samples in both plots. Number (0-30) in the legend represents the day of sampling. The middle panel shows the (B) trends of top 15 phylum. (C) The scaled heat-map showing the of number of overexpressed differential genes in different pathways for the two sediment types. Please refer to Figure S2 for the heatmap of individual functional gene abundance trends.

Specific taxa of primary producers, such as cyanobacteria and gemmatimonadetes phyla, formed the first ecological group, whose abundance increased from about 20 to 50% between day 0 and day 1 in the acclimation autotrophy phase (days 0 - 3). The functional genes that encoded for the pathways of photosynthesis and nitrogen metabolism were largely mapped to the members of cyanobacteria (Figure 5). Their higher abundance in the industrial land use compared with the residential land use is in agreement with a previous study ^5^ and is related to the lower levels of antibiotics in industrial areas ^1^. The heterotrophic ecological group in this phase was mostly dominated by proteobacteria, which encoded functional genes for alternate sources of carbon, such as the products of organic degradation.

**Figure 5:**
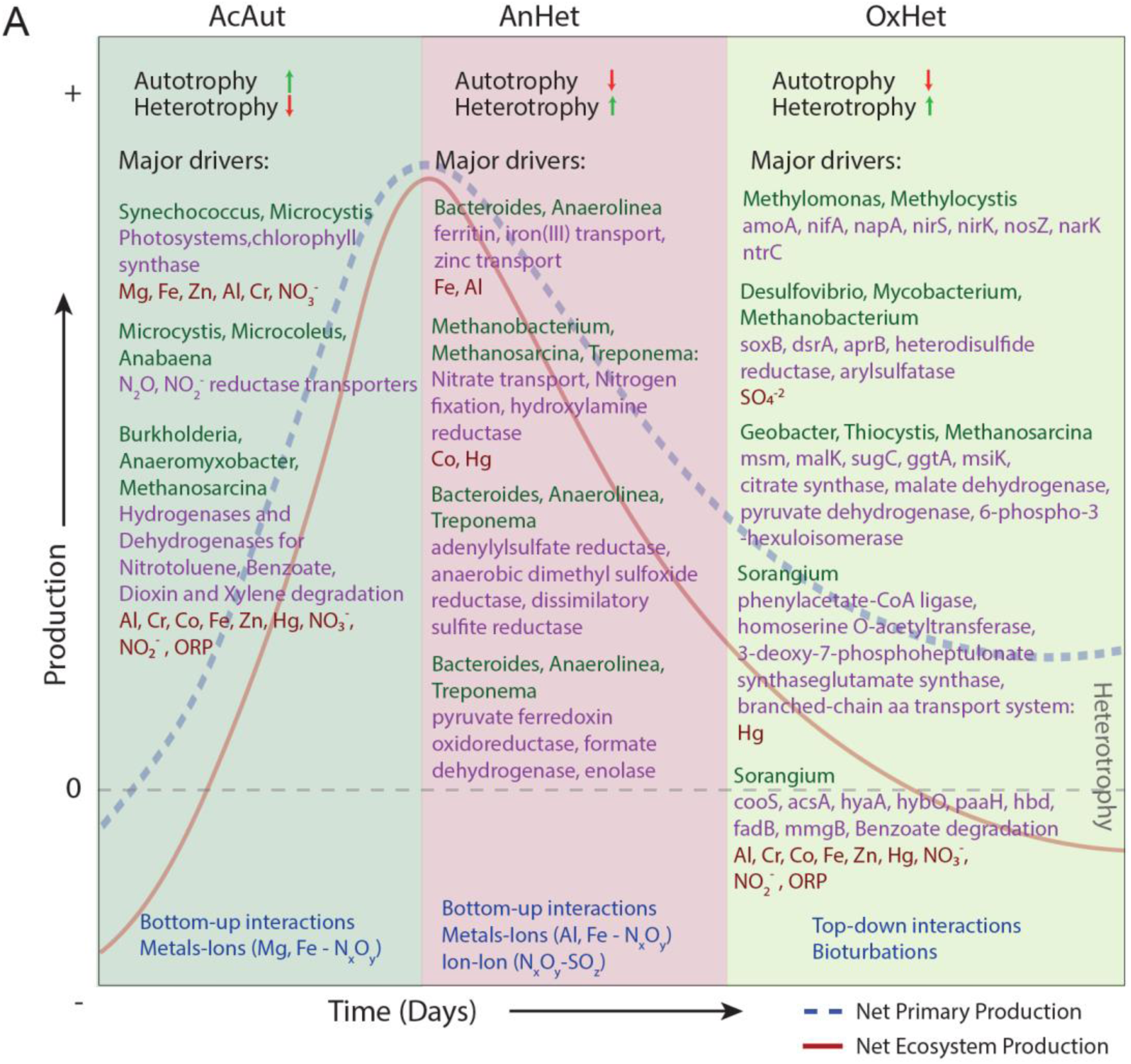
Overall summary model depicting the major drivers of net primary productivity (NPP), net ecosystem productivity (NEP) and heterotrophy during the temporal-shift phases in sedimentary microbial ecosystem. Green and red arrows represent the upward and downward trend of microbial process/functions genes, respectively in different temporal shifts stages. The heterotrophy is defined here as the difference between NPP and NEP. The major taxonomic groups at genus level, functional genes in respective pathways and the environmental parameters driving each phase are shown below the model.

The anoxic heterotrophy phase was characterized by an increase in biomass, a reduction in primary producers (∼10%), DO and ORP, and a concordant increase in heterotrophs (Figure 4B). The reduction in photosynthesis-related genes (Figure 4C) corroborated the decreasing trend in expressed functional genes for phototropic-based carbon fixation. Interestingly, the nitrogen metabolism functional guild, which was driven by cyanobacteria in the oxygen-rich acclimation autotrophy phase, was taken over by Archaea and Spirochaetes under the reducing conditions of the anoxic heterotrophy phase (Figure 5). The composition of the heterotrophic functional guild also changed significantly, with an increase in Bacteroidetes, Chloroflexi and proteobacteria species that were different from the acclimation autotrophy phase. Significant functional overlap among these community members was observed in this phase; for example, the functional genes of metal transport and metabolism of sulfur and simple sugars were mapped primarily to *Bacteroides* and *Anaerolinea* (Figure 5). Similarly, *Treponema* co-expressed the functional genes of sulfur, nitrogen and simple sugar metabolism (Figure 5).

The oxic heterotrophy phase showed a significant increase in DO and ORP values, compared with the preceding phase. Although phylum level community composition in this phase appeared similar to that of the anoxic heterotrophy phase, there were significant differences at the level of species and functional guild (Table 1A, S1 and 1B). The biomass increased in the initial days of this phase coincided with a complete shift in the active functional guilds from the previous phase (Figure 4C and 5). The functional overlap that was observed in the anoxic heterotrophy phase was not observed in this phase. On the contrary, potential resource niche speciation was observed, with different functional guilds mapping to different sets of microbial community members (Figure 5). Overall, a photosynthetic guild was very poorly represented, sulfur and nitrogen oxidation were active, and simple sugar metabolism and organic degradation was driven by proteobacterial species (Figure 5).

Although, the trends of microbial community shifts were conserved in the two land-use types, the chronic pressure from land-use types influenced the microbial community composition. For instance, representation of the proteobacteria phylum did not differ significantly in abundance over time. However, their representation was significantly different between the two land-use types at species level at all timepoints (Table S2). Acidobacteria, with the potential to degrade plant-, fungus- and insect-derived organic matter and utilize a broad range of complex carbon substrates were more abundant in industrial land-use zones. However, methanogens such as euryarchaeota and members of phyla such as Bacteroidetes, Chlorobi and Firmicutes were differentially abundant in the two land-use types. Most unassigned differential taxonomic groups at genus level were abundant in the residential areas (Table S2), which not only explains the higher diversity of microbial communities in the sediments of residential land-use types, but also points to the TruCS in residential areas as potential regions to discover novel microbial processes.

While the dynamics of microbial communities were consistent between the land-use types, their composition varied significantly. PERMANOVA analysis of the active (Table 1A) and total community (Table S1) showed significant differences between the land-use types and between days for both datasets. The differences in the transcriptome levels of SedMICs in the two land-use types and over time were statistically supported by the PERMANOVA analysis. There were significant differences in gene expression between the two land-use types. Over time, such differences were significant between days 0 and 12, days 1 and 30 and days 12 and 30 in both land-use types (Table 1B). Therefore, these days 0, 12 and 30 can be representative days of the three phases of ecosystem shifts after the rain-perturbation in TrUCS.

### Specific metal-ion combinations are associated with distinct microbes and their processes

Specific ecosystem descriptors showed interactions with the functional guilds from certain microbial community members of the two land-use types. Two ecologically different functional groups emerged from significant and strong correlations between particular combinations of metals and nutrients with different groups of phyla (Table 2A) and their associated functional genes (Table 2B). These groups formed the basis of a metal-nutrient-microbial axis of the ecosystem functioning of TrUCs. The first metal-nutrients group comprised magnesium-nitrates that significantly correlated with the ecological consortium formed by cyanobacteria, gemmatimonadetes and nitrospirae. These taxonomic groups are known for their photosynthetic abilities and nitrogen metabolism. Interestingly, strong and significant correlations of magnesium (0.7), iron and zinc with cyanobacteria were observed, together with genes from photosynthetic pathways. The second metal-nutrients group of sulfates, nitrates and metals including aluminum (Al), cobalt (Co), lead (Pb) and mercury (Hg), correlated significantly with an ecological consortium of phyla including chlorobi, CDOP11, chloroflexi and euryarchaeota. These phyla contain members capable of sulfur metabolism and degradation of organics. Similarly, genes from organic remediation pathways correlated significantly with nitrates and metals, such as aluminum (Al), cobalt (Co), lead (Pb) and mercury (Hg).

**Table 2:**
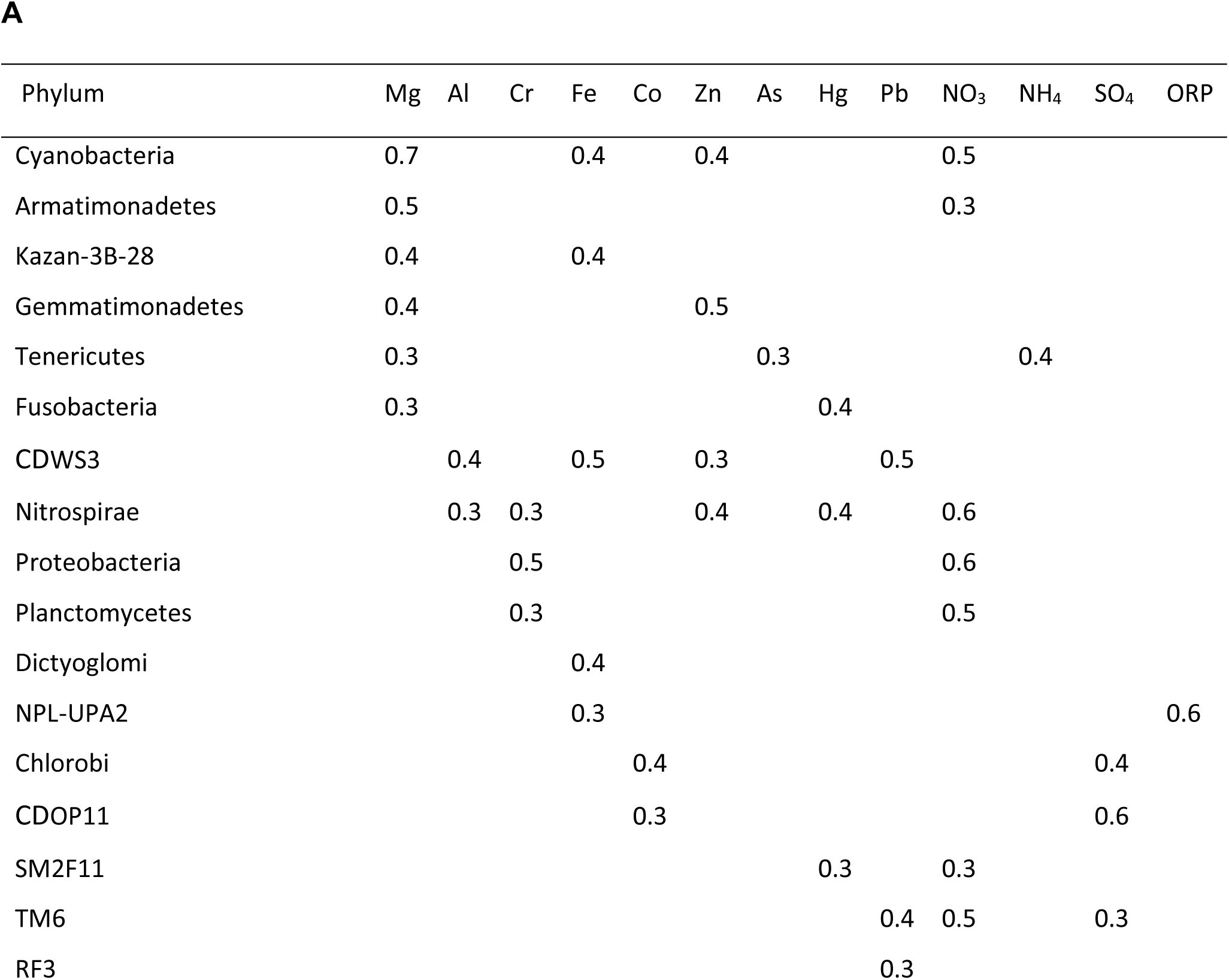
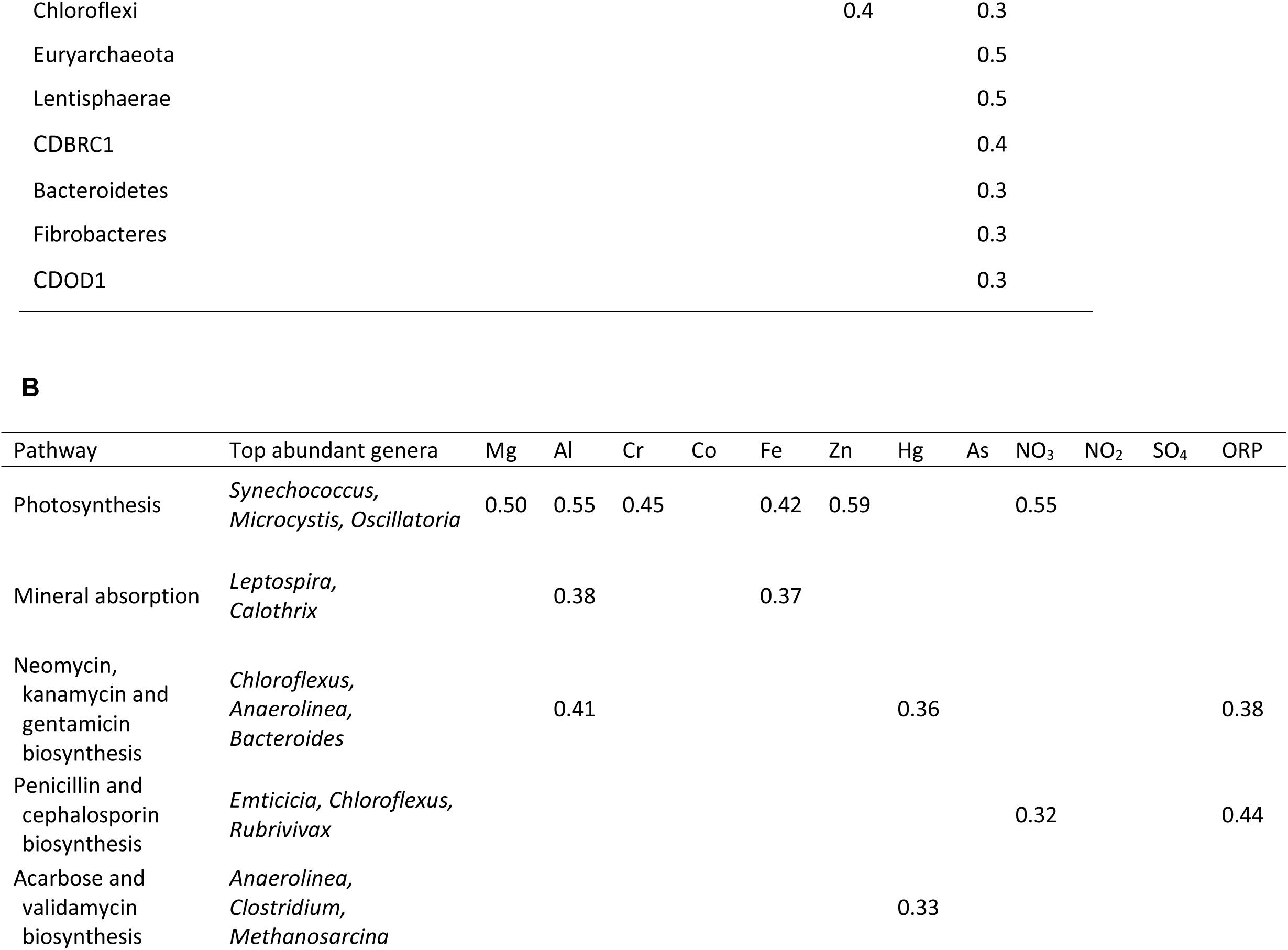
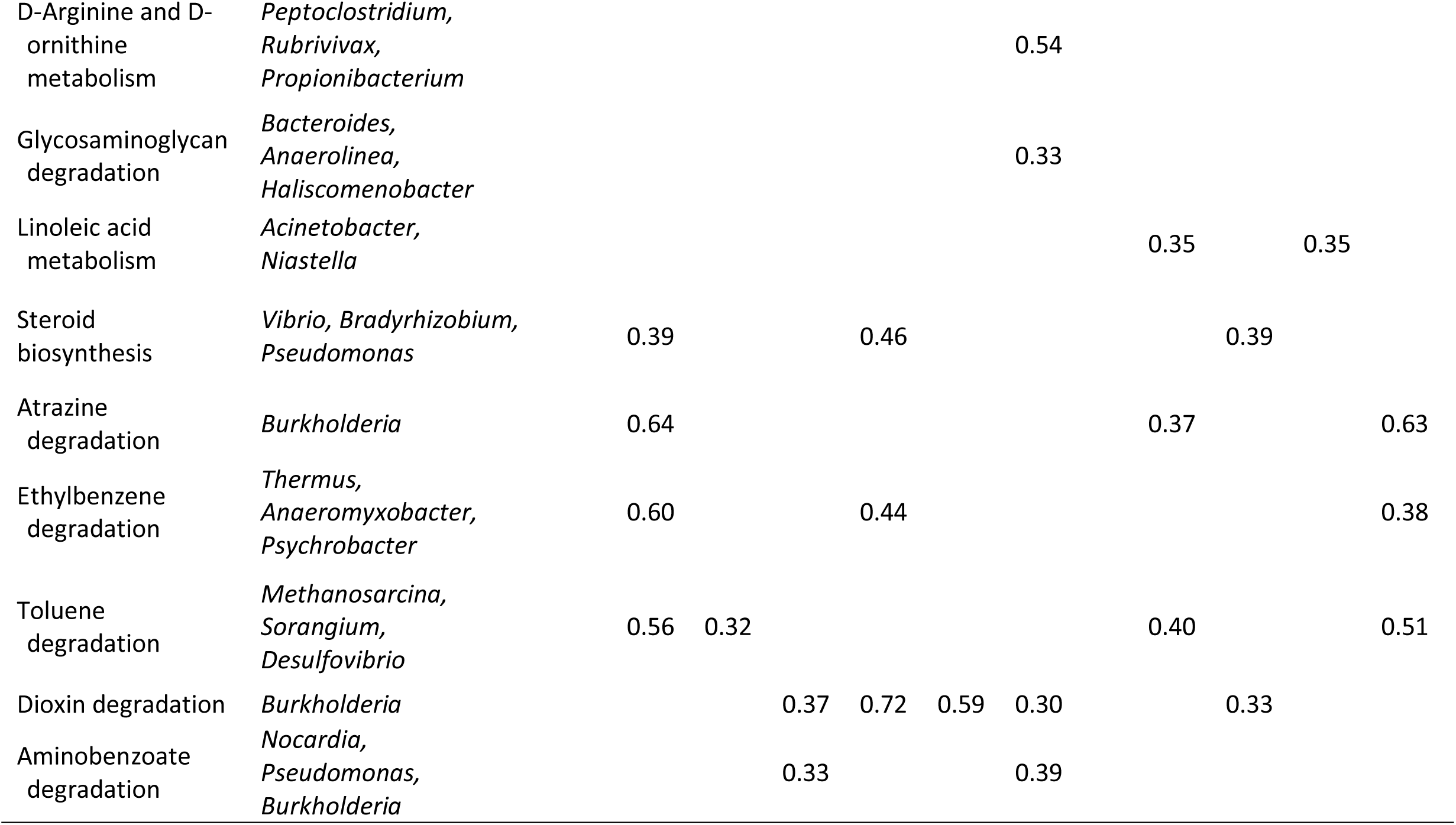
Spearman rank correlation between ecosystem descriptors and (A) microbial taxa aggregated at phylum level and (B) functional genes aggregated at pathways level. Significant correlations (*p* < 0.05) above 0.3 are shown here.

## Discussion

The structure and function of microbial communities in well-managed and controlled canal ecosystems have been studied in the context of specific physicochemical stressors ^1^. In this study the associations of ecosystem descriptors and microbial communities were identified at a catchment scale in a spatial analysis ^1^. Here, we uncover the major ecosystem trends of SedMICs and associated ecosystem processes in TrUCS using an ecogenomics framework ^1^, which is augmented by integrated multi-omics approaches. The microbial ecology patterns and processes identified here are in the context of a natural intermittent disturbance (rain) and its influences on SedMICs in two different anthropological chronic disturbance (land-use) regimes. We have disentangled the metal-nutrient axis, identified specific associations with the microbial pathways and presented these in a microbial ecology framework of primary productivity and heterotrophy. The association of metals and ions has been well-established for specific microbial species ^17, 6, 1, 18, 19^. However, the increased complexity present at the level of specific metal-nutrient associations, together with their role in ecological shifts in microbiome functional guilds, presents an innovative way of investigating temporal changes in sediments using ecosystem descriptors.

Here we present four major outcomes regarding shifts in SedMICs and associated ecosystem processes. Firstly, the shifts in the sedimentary ecosystem are rapid (hours to days), compared with the macro-ecosystems (years) ^9^. Secondly, distinct phases of the temporal shifts are clearly visible at the level of geochemical variables, community composition and ecosystem functional guilds. SedMICs that differ in composition in different land-use types have conserved intrinsic patterns of shifts in community functions. The rapid changes in the sedimentary ecosystem, and their distinct timing, would enable urban water resource managers to plan and implement interventions to maintain the ecosystem primary productivity and diversity. Thirdly, metals and nutrients are significantly associated with, and influence ecosystem functioning through their involvement in energy capture from photosynthesis to energy generation, such as linked nitrogen-sulfur transformations and energy utilization for organic degradation. Lastly, the influence of sedimentary ecosystem functions is clearly reflected by the water quality parameters.

The dynamics of the sedimentary ecosystem of TrUCS are explained by the integration of multi-omics data through statistics and modeling (Figure 5). The first major outcome is the rapid temporal shifts in the ecosystem functioning of net primary productivity (NPP) and heterotrophy, driven by the biogeochemistry and SedMIC functions. These dynamic shifts occurred within four weeks of a disturbance in both residential and industrial land-use types and are consistent with the phenomenon of rapid microbial processes in a tropical climate ^20–22^. Compared to macro-ecosystems, microbial generation time is short and in sedimentary ecosystems functional shifts occur in timescales of days compared to, for example, decades for forest ecosystems ^13^.

The second major outcome, the ecological shifts, was deduced based on clustering patterns of ecosystem descriptor data. Similar patterns were observed in microbiota composition and function data. The shifts in the functional guilds, sediment geochemical profiles and their direct influence on water quality are spread over three distinct phases, namely, acclimation autotrophy, anoxic heterotrophy and oxic heterotrophy. The complete functional regime shifts in the three phases of urban freshwater canals indicates that changes in SedMICs and functional turnover rates occurr within a few days, and hence communities are very active after the rain event.

In the acclimation autotrophy phase, primary productivity and nitrogen reduction dominate the community functions led by cyanobacteria, which provide the simple sugars for community survival after a pulse disturbance of rain. Heterotrophy involving organic degradation by microorganisms is also observed, and enabled by the seed microbial community entering the canal with rain water. The strong correlations (r > 0.5, *p* < 0.05) of Mg, Fe and nitrates with primary producers and their major functions, such as cyanobacteria and photosynthesis, clearly demonstrate metal-nutrient assisted primary productivity in a sedimentary microbial ecosystem. Further, this correlation indicates that energy capture, production of simple sugars and nitrogen reduction are as essential to the sedimentary ecosystem after perturbation as they are for ecosystems inhabited by macrorganisms, such as forests ^13^. Similar assessments of increased carbon and nitrogen fixation were observed in the microbial communities from a hypersaline Bahamian lagoon after a disturbance by Hurricane Frances ^23^. While the autotrophs are engaged with primary productivity, the heterotrophs, such as, *Burkholderia, Anaeromyxobacter* and *Methanosarcina* sustain the microbial community through the utilization of alternative sources of carbon, such as nitrotoluene, benzoate, dioxin and xylene by increased expression of functional genes like hydrogenases and dehydrogenases. This heterotrophy guild is strongly correlated with the second metal-nutrient set comprising of Al, Fe and sulfates, delineating the different drivers of heterotrophy fueled by organic degradation and availability of simple sugars. It shows that utilization of alternative sources of carbon through organic degradation requires specific metal-nutrient intermediaries. Further, incoming microbial communities appear well adapted with a functional guild capable of organic degradation ready to utilize the alternate sources of carbon in a perturbed environment where resources, such as simple sugars, are limited.

The outcomes of nitrogen reduction in the acclimation autotrophy phase and metal sequestration appears in the anoxic heterotrophy phase. For example, anoxic heterotrophy phase shows high levels of ammonium and a reduction in the levels of metals in the loosely attached fraction in sediments. Further verification of the role of metals in sedimentary ecosystem functioning is provided by the ability of the microbial community to sequester metals. This function is evidenced by the expression of genes for mineral transport, such as ferritin, iron(III) and zinc in *Bacteroides* and *Anaerolinea* spp. In a reduced environment that starts to develop after day 3, metal speciation begins shifting to the lower oxidation states and metals thus form composites with carbonates, sulfides and humic substances ^24^. These composites appear to be bioavailable to the microbial communities ^25^. With an abundance of simple sugars in the anoxic heterotrophy phase, a heterotrophic guild displaces the photosynthetic guild. Similar observations were reported in intermittent waterways, where anoxic conditions were associated with increased heterotrophy^26^. Within a seven-day period the microbial community of the anoxic heterotrophy phase shifts completely in terms of compositional and functional attributes, signifying a rapid community transformation. Heterotrophs utilize carbon and nutrients made available from the microbial communities of the acclimation autotrophy phase, rather than relying on the external sources. The simple sugars provided by the microbial communities of the acclimation autotrophy phase are utilized by the succeeding subsequent microbial communities. This is evidenced by a shift community structure and function following the acclimation autotrophy phase, with abundant differences in heterotrophic taxa, including members of the *Bacteroides, Anaerolinea* and *Treponem*a, and increased expression of functional genes such as pyruvate ferredoxin oxidoreductase, formate and dehydrogenase, and enolase.

Such a shift in community composition in the anoxic heterotrophy phase and an increase in heterotrophic bacteria and concordant increase in the utilization of simple sugars is comparable to the increase in heterotrophy in ecosystems defined by macroorganisms ^13^. Together with the reduction in cyanobacterial abundance, the available oxygen levels drop, creating a reducing environment in the ecosystem. In such conditions, nitrates could be utilized as the terminal electron acceptor (nitrate respiration) ^27^ for the generation of metabolic energy. The expression of nitrogen reduction and metal transport genes that was initiated in this phase (anoxic heterotrophy), continues to increase, however, through different species than in the acclimation autotrophy phase, such as, *Methanobacterium, Methanosarcina* and *Treponema.* As a result, ammonium levels increase and nitrate levels drop during the anoxic heterotrophy phase, both in sediments and in bulk water. The significant correlation of Co and Hg with amino-benzene degradation pathways could partially explain the increase in ammonium and these metals during the anoxic heterotrophy phase.

Interestingly, in the oxic heterotrophy phase, the cyanobacteria do not recover. The oxic heterotrophy phase continues to maintain the high expressions of simple sugar and amino acid metabolism pathways, along with the organic degradation pathways, although the community members in this phase differed from those of acclimation autotrophy and anoxic heterotrophy phases. In addition to this, low levels of nitrates and high levels of ammonium would have stimulated the ammonium assimilation, as reflected by the increase in biomass. Thus, this phase is likely to be formed by three major guilds – nitrogen assimilation, sulfur metabolism and heterotrophic – which are driven by different sets of microbial community members across phases. The assimilation of nitrogen is taken over by species such as *Methylomonas and Methylocystis.* Similarly, the sulfate levels start to build up through actions of another set of community members, such as *Desulfovibrio* and *Mycobacterium*. This coincides with the increase in cell count, amino acid metabolism and expression of protein synthesis genes, indicating the assimilation of nitrogen is coupled with sulfur oxidation ^28^. The top-down bioturbation effects are also apparent in the latter half of the experiment and seem to have assisted the oxidation of nitrogen and sulfur by establishing the oxidative environment in this phase. The experimental design of this study limits the immigration of sedimentary microbial community members, which explains the collapse of microbial communities at the end of oxic heterotrophy phase.

The compositional and functional shifts from primary productivity-dominant to heterotrophy-dominant SedMICs also indicate a transformation of the SedMICs ecosystem from a carbon sink to carbon source. A controlled and informed release of catchment microbial communities (with cyanobacterial population) and intermittent dredging of canal sediments could potentially restrict the anoxic environment and thus the conversion of the SedMICs ecosystem from a carbon sink to a carbon source. In addition to primary productivity, an important ecosystem service of degradation on organic contaminants, which remains high in the first and third oxic phases along with higher biomass, could be maintained using the same approach of intermittent sediment dredging.

Characteristic temporal phases of microbial communities, their functions and the resulting outcome on the biogeochemical levels were consistent in the residential and industrial land-use types. This shows that the urban sedimentary ecosystems have conserved or similar functional guilds, which are driven by different taxonomic entities or strains, suggesting functional deterministic processes may influence community functions (along with stochastic processes), at least in the context of this study. It further suggests the direct influence of sedimentary microbial processes on surface water quality in aquatic systems, irrespective of land-use influence from the catchment. Such a conserved response can inform and direct urban waterways management by adopting relevant targeted approaches.

In conclusion, this study demonstrates how ecological theories that explain post-disturbance dynamics in macro-organism ecosystems, can be applied to smaller sedimentary microbial ecosystems, despite much shorter time-scales in the latter. Understanding system behavior from a molecular ecology perspective provides highly detailed mechanistic insights into the top-down and bottom-up influences that drive the system dynamics, especially for healthy ecosystem functioning. Such a framework can enable rationally designed monitoring and management for an enhancing self-cleaning capacity of canal systems, and can be extended to highly dynamic ecosystems that have a strong input from microbial processes.

## Methods

### Mesocosm flumes

Mesocosm flumes (Figure 1A) were fabricated using 1.8-cm-thick acrylic glass sheets. Each flume was 235.1 cm long (internal length: 231.5 cm), 18.6 cm wide (internal width: 15 cm) and 16.8 cm high (internal height: 15 cm). A space reserved for sediment was provided in the centre along the entire length of the flume. A 10-cm-long area at the extreme ends of the flumes was isolated using 0.5-cm-thick partitions with uniformly distributed holes that were 5 mm deep in the top portion (10 cm from the top) and 2 mm deep in the bottom portion (5 cm from the bottom). This arrangement smoothed the flow of recirculating water and prevented the packed sediment bed from being washed away. Each flume was fitted with its own recirculation and replacement circuit. The recirculation circuit included water outflow, sink, connecting polyvinyl chloride (PVC) pipes (32 mm), a centrifugal pump (Eheim 5000, 41.5 - 83.3 L/min) and gate valves. Centrifugal pumps were run at full workload and excess water on the pressure side was bypassed to the sink to avoid any resistance buildup on the pump. Only the required amount of water was allowed to flow towards the flume inlet. The replacement circuit included a replacement water tank (PVC, 150 L), a peristaltic pump (MasterFlex L/S® Easy-Load® II), connector silicon tubes (5 mm), PVC piping and two-stop pump tubing (silicone platinum-cured tubes: MasterFlex L/S® 16). To maintain the flow across fresh water inlets and outlets, a single peristaltic pump was used for incoming as well as outgoing water. Fresh water was stored in separate tanks for each flume. Fresh water was filtered through a mesh (≈1 mm) to restrict the debris from entering the flume system. The whole set-up was kept on an elevated platform shared by two flumes. PVC sheets were raised between the flumes to prevent cross contamination. Flumes were covered with aluminum foil to reduce light penetration and discourage algal growth.

### Hydrodynamic settings

The flumes were operated at continuous recirculation and replacement of water with constant flow for 30 days. The recirculation was maintained at ≈6.3 L/min (±5%), in laminar flow with mean bulk velocity of water above the sediments ≈1 cm/s (±5%) at all times during the experiment. In the present study, to calculate Reynolds number for rectangular open flow channel, hydraulic radius was first calculated as:

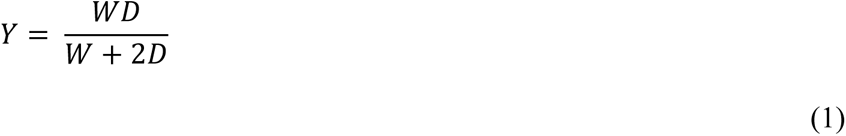

Where *Y* is the hydraulic radius (cm), *W* is width of flume (cm) and *D* is water head over sediments (cm). The Reynolds number was then calculated using following formula:

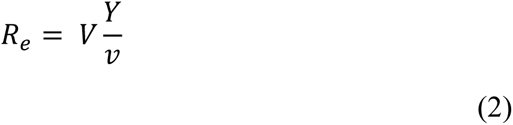

Where *R_e_* is Reynolds number, *V* is the bulk velocity of water (cm/s) and is kinematic viscosity (cm^2^/s). The outflow was kept at 10 cm from the bottom (inner surface) of the flume to maintain a constant water head of 7cm above the sediment bed. Outflowing water was captured in the collection tank and was pumped back upstream in the flumes for recirculation. Flow of fresh water was adjusted to replace the full flume volume (≈24 L) in about 13 h. Fresh water was continuously injected from the replacement tank to the collection tank while it was removed from the downstream end of the flumes at a constant flow rate of 32 mL/min (Figure 1A). Fresh water was brought from the same locations and the replacement tank was replenished every 48 h.

### Experimental design

With the objective to cover the full temporal shifts, a 30-day experiment was designed for this study with ten sampling time-points. The experimental design had three factors (Figure 1B). Sediment type and time were fixed factors and flume number was a random factor nested in sediment type. Two sediment types, residential and industrial, were studied during the experiment. Each sediment type was run in three parallel replicated experiments and each experimental replicate was sampled three times as technical replicates. The experiment was randomized at each level (unless otherwise stated) and conducted in a semi-open environment with protection from rain.

### Seeding of flumes

Sediments and water were collected from two sites, which are representative of residential and industrial land-use types in the Ulu Pandan catchment of Singapore ^1^. The industrial site is located in the Jurong east area (1.332133, 103.749433) and the residential site in the Clementi area (1.318195, 103.771985). Sediment and water were collected from both sites on the morning of 23^rd^ November 2012 after a rain event. The duration between rain event and sediment/water collection was between 6 and 8 h.

Freshly settled top layers of sediments were carefully collected and transferred into clean buckets. Water was collected from the same site in pre-cleaned 25-L carboys and immediately transferred to the experimental site. The sediments were homogenized and cleared of coarse wood and leaf debris before spreading into the flumes to the height of 3 cm. Ceramic cups, made of natural clay were equilibrated with canal water for 15 minutes before they were buried in the top 1 cm layer of sediments. Water from each respective source was added to flumes to start the recirculation circuit. Replacement tanks were also filled with the same water and the replacement circuit was also switched on. The flumes were monitored daily for water flow and water quality indicators. The replacement tanks were filled with fresh water from respective locations every alternate day.

### Sampling

The sampling was divided into two parts: sampling on time points and daily monitoring (Figure 1C). Sediment samples were collected on the time-points for meta-omics and meta-environment parameter measurements. We performed 16S amplicon sequencing and metatranscriptomics. Meta-environment measurements assessed metals, nutrients (ammonium, nitrates, nitrites, sulfates and phosphates) and biomass (cell-counts). Relevant methods are described in more detail below (*Nutrient and metal analyses*). Ceramic cups with 2.5 cm internal diameter and 0.5 cm depth were buried 1 cm in the sediment bed of the flumes for sediment sampling. The spacing between the ceramic cups was about 3 cm from each other and from the side walls. Samples were collected in triplicate from random locations within the flume to reduce possible linear trend due to linear water flow. To avoid sample artifacts at the extreme ends, samples were not taken in the outer-most 10 cm region of the both ends of the sediment bed. Water samples were also analyzed for metals and nutrients during the time-points. For the daily monitoring, water samples were tested for physicochemical parameters such as temperature, pH, conductivity, salinity, dissolved oxygen, oxidation-reduction potential (ORP) using YSI 556 multi-probe system and 650MDS data recorder from Xylem Inc.

Water samples were collected in clean 15 mL Falcon™ tubes and kept on dry-ice during transportation. For sediment sampling, buried ceramic cups were carefully taken out without disturbing the nearby sediments and the contents were immediately transferred to clean 50 mL Falcon™ tubes filled with pre-chilled buffer and stored on dry-ice. The samples were transported immediately to the laboratory for further processing. Water samples were stored in freezer (−20°C) till analysis. Sediment samples were processed on the same day for DNA extraction and stored in −80°C until processing for RNA extraction.

### Nutrient and metal analyses

#### Samples preparation

Metals and nutrients loosely attached to sediments, soluble in pore water and bioavailable to the microorganisms were measured. The protocol to extract the water extractable fraction of metals and nutrients was adapted from previous reported methods ^29, 30^ with slight modifications. Extractions of 1 g of sediment samples were performed with 20 mL of Milli-Q water (18.2 MΩ·cm) added to the samples. The samples were shaken at 200 rpm for 16 h at room temperature to detach the sediment surface-associated inorganic compounds. Extracted inorganic compounds in the supernatants were filtered using 0.2 µm filter. The filters were flushed first with acidified Milli-Q water (pH 3) and then with neutral pH water prior to sample filtration. Nutrient ions were tested from 10 mL of supernatant using Prominence HIC-SP IC dual system for anion and cation simultaneous analysis system from Shimadzu. Remaining 10 mL of water was acidified with 500 µL of analytical grade nitric acid for metals analysis using ICP-MS (Agilent 7700).

#### Nutrient quantification

Commercially available certified solutions from Merck, Darmstadt, Germany were used to prepare standard curves for measuring nutrients. Samples were diluted 10x and transferred to clean ion chromatography (IC) vials. One milliliter of sample aliquot was analyzed using Shimadzu Prominence Ion Chromatography dual system (Shimadzu, Japan). The sample injection was performed using 500 µL sample into the ion chromatography machine through auto-sampler (Shimadzu, Japan). Anions were analyzed using column shim-pack IC-SA2 and cations with shim-pack IC-C3. Sodium bicarbonate (12 mM) and sodium bicarbonate (0.6 mM) in water (MERCK, Germany) were used as solvents for anions and injected at a flow-rate of 1 mL/min. The solvent used for cations was 2.5 mM oxalic acid in water (Sigma Aldrich, USA), which was injected at a flow-rate of 0.95 mL/min. Areas of detected peak were integrated with the retention time (RT) of ions of interest using Shimadzu LabSolution software V5.51. Concentrations of ions of interest were calculated from the standard curve within its linearity range.

#### Metal quantification

Samples were acidified with 500 µL of analytical grade nitric acid and diluted 10x before transferring into clean 5 mL ICP-MS vials. The introduction of samples to (inductively coupled plasma mass spectrometry) ICP-MS was automated via an auto-sampler attached to the instrument. The ICP-MS instrument was tuned for helium in spectrum mode to detect multiple elements in single run. Sample uptake time was set to 20 s, while stabilization time was set at 30 s. Washout procedures followed the run of standards and at an interval of 10 samples. The procedure included two 40 s washes with ultraclean Milli-Q water separated by a 2% HNO_3_ wash for 30 s. Randomly picked standard and blanks were tested after every 15 samples and 20 samples, respectively, for quality control. Standard curves were prepared from using ICP-MS complete standard - IV-ICP-MS-71A in HNO_3_ and mercury for ICP-MS in HCl from Inorganic Ventures Virginia USA, and run for each set of samples.

### Cell counting

For cell counting, the protocols were adapted from previously reported studies ^31, 32^. Sediment samples were dewatered and 0.2 g placed in 2mL Eppendorf tubes. Samples were sonicated at 37 kHhz, 320 W, for 15 min at 35°C in 1.5 mL 10% methanol (90% filtered phosphate buffer saline (PBS), pH 7.4), then placed on ice for 5 min. Sonication and cooling on ice was repeated. The samples were then centrifuged at 200 g for 2 min to remove unwanted sediment particles. Equal volumes of supernatant with dislodged cells were transferred into two Eppendorf tubes (0.75 mL each). The contents of each tube was centrifuged at 8000 g for 5 min to obtain a cell pellet, and the supernatant removed. In one tube, 0.2 mL of 4% paraformaldehyde (PFA) was added for fixing Gram-negative bacteria. In the other tube 0.2 mL of 100% ethanol was added to fix Gram-positive bacteria. Samples were incubated overnight at 4 °C. Cell pellets were again obtained by centrifuging samples at 8000 g for 5 min, then was washed twice with 0.2 mL filter-sterilized PBS and re-suspended in 0.5 mL of equal volumes of 100% ethanol and PBS. The sample (0.1 mL) was then mixed with 0.5 µL SYBR Green and incubated in the dark for 30 min. The samples were again centrifuged at 8000 g for 5 min, washed, re-suspended in 0.5 mL PBS and transferred to flow cytometry tubes, for cell counting. Separate gating boundaries for PFA and ethanol fixation conditions were determined with different stain concentrations (0-5 µL of 100x working solution of SYBR Green), with and without (negative control) sediments. Samples from day 30 were used to confirm the gating boundaries. The threshold was set at 800 on PF-1 (Green fluorescent) under both fixation conditions.

### Nucleic acids extraction

Genomic DNA was extracted from the samples immediately after collection. Sediments were dewatered and DNA was extracted by a combination of mechanical, chemical and thermal lysis and chloroform-isoamyl alcohol purification using protocol described by Zhou et al (1996) ^33^. Briefly, 5 g of dewatered sediments were mixed with DNA stabilization buffer described elsewhere ^33^. Mixture was transferred to pestle and mortar, pre-chilled with liquid-nitrogen. The samples were ground and freeze-thawed thrice by flooding with liquid-nitrogen. The contents were then transferred to lysis buffer ^33^ and heated to 65 °C for 3 min. Proteinase K (20 IU) was added to the mixture and contents were heated again for 2 h, mixing the samples every 20 min. Samples were then centrifuged at 10,000 g for 20 min at room temperature prior to supernatant being collected in a clean tube without disturbing the layer between liquid and sediment phase. The step was repeated to further remove impurities. The contents were added with equal volume of 1:24 isoamyl-alcohol:chloroform and gently shaken for 5 min before centrifuging at 14,000 g. The resulting supernatant was collected without disturbing the impurities settled at the interphase. This step was repeated twice. The supernatant was then mixed with 0.6 volume of absolute iso-propanol and incubated overnight in −80 °C. The samples were thawed at room temperature and centrifuged at 20,000 g for 20 min, with the resulting DNA pellet recovered and re-suspended in nuclease-free water. Removal of co-precipitated humic substances was achieved by OneStep™ PCR Inhibitor Removal Kit from Zymo Research Corporation (USA) following manufacturer’s protocol. DNA was suspended in nuclease free water and quantified using Qubit®2.0 fluorometer (Life Technologies, USA).

Total RNA extraction was carried out using RNA PowerSoil® Total RNA kit (MOBIO, USA) using 2 g sediments, following the manufacturer’s protocol. Extracts were quantified using Qubit®2.0 fluorometer (Life Technologies, USA). DNA contamination was removed using RTS DNAse kit (MOBIO, USA), following the manufacturer’s protocol. RNA quality and quantity were assessed using Agilent 2100 Bioanalyzer (Agilent, USA).

### Sequencing

#### Transcriptomics

Days 0, 1, 4, 8, 12 and 30 total-RNA-seq of sediments from one sample per flume of residential and industrial sediment types were performed using Illumina HiSeq2500 system. This resulted in metatranscriptome data for 36 samples (two sediment types × three samples per sediment type × six time-points). The raw cDNA read data was quality trimmed using cutadapt-1.5 (using default parameters except –overlap 10 -m 30) and sand processed to remove reads likely to originate from rRNA genes by filtering against Ribosomal Database ^34^. Subsequently, all remaining reads were subjected to an homology search against the RefSeq non-redundant proteins database ^35^ (Sep 2015) using diamond v0.7.9.58 ^36^. The diamond output was processed using the command-line version of MEGAN ^37^ (blast2lac in megan_ultimate_6_1_18 suite) and then further reduced using custom scripts in R to generate KEGG (KO identifiers) and taxa assignments. The files were processed to generate a KO-by-sample matrix of raw read counts. This KO-by-sample matrix was further normalized using R/DESeq ^38^ to standardize for inter-sample differences in total read counts using a variance stabilization and calibration method ^39^. In cases where zero counts were detected the element of this matrix was coded as “NA”. Active microbial community members were identified using v5 region of 16S rRNA transcripts. The transcripts were screened and annotated using NCBI database search for understanding changes in the active microbial community.

#### Phylogenetic classifications

Following the DNA extraction and purification, v1 – v6 regions of the 16S rRNA gene were amplified for one replicate from each flume for all time points using universal primer sets for microbial communities ^40^. The amplicons were attached with barcodes specific to the samples and sequenced on Illumina MiSeq2500. The phylogenetic classifications of the resulting 16S rRNA SSU amplicon sequences was performed using the QIIME ^41^. Briefly, paired-end reads from sequencing data were first filtered for quality, using cutadapt ^42^ and then joined using FLASH ^43^. The joined reads were filtered again with more stringent parameters and then demultiplexed. High-quality reads were used for operational taxonomic units (OTU) picking using the USEARCH algorithm ^44^. After OTU-picking, the taxonomy was assigned to the representative sequence of each OTU cluster. The most recent version of SILVA database ^45^ was used for the taxonomy assignment. Data in the OTU table thus created were normalized for sequencing depth to allow comparison between samples.

### Statistical analysis

All statistical analyses and graphs were produced in R ^46^, except for PERMANOVA analysis and the distance-based linear modeling (DistLM), which were done in PRIMER v6 (PRIMER-E, UK) ^47^. Ecosystem descriptors, cell count, and daily monitoring datasets were pre-processed as follows. Blanks were subtracted to account for the blank’s presence in the metal’s measurement. The quality control-robust LOESS signal correction (QC-RLSC) was used to correct for and minimize the impact of analytical variation in the physical-chemical variables, which could be due to changes in signal response or instrument calibration. This algorithm incorporates a locally estimated scatterplot smoothing function (LOESS) with a span of 0.75 that is fitted to the QC data with respect to the order of extraction (independent variable). We used the LOESS function from R. A correction curve for the complete data matrix is then interpolated along the order of extraction, to which the total data set for that feature is normalized. The described method reduces the technical variation observed ^48, 49^. Drift in the signal for each metal was corrected for by observing the change in the signal for the same metal across standardized quality control samples (50 ppb for magnesium, 2 ppb for mercury, and 20 ppb for all other metals) and correcting the samples for any temporal shift across extraction times in the signal related to the observed quality control samples.

Unless examining a relative measure such as relative abundance, the regularized logarithm transformation (rlog) from the R package DESeq2 was applied to the operational taxonomic units (OTUs) and functional genes to render them homoscedastic and to account for differences in sequencing depths between samples ^38, 50, 51^.

Contour plots ^52^ of metal, nutrient, cell count, & daily ecosystem descriptors over time (days after rain) were created using scaled values of datasets over time (mean = 0, standard deviation = 1). The ecosystem descriptors were clustered into five groups with similar means over time using hierarchical k-means clustering (hkmeans) from the R package factoextra (https://cran.r-project.org/web/packages/factoextra/index.html). The hkmeans algorithm is designed to ameliorate k-means’ sensitivity to the initial centroid locations. Hierarchical clustering, with a complete clustering method and Euclidean distance measure, was used to determine the initial centroid locations. K-means clustering was then performed using these initial values ^53^. The scaled values of the ecosystem descriptors were averaged within the clusters they belong, smoothed using locally estimated scatterplot smoothing (LOESS, span = 0.75), and plotted.

The Spearman’s correlations between log_2_ transformed ecosystem descriptors data and rlog transformed microbial communities composition aggregated by phylum and rlog transformed functional genes aggregated by pathway are presented showing only those correlations that are significant (p < 0.05) and above 0.03. Bray-Curtis dissimilarity-based principal coordinates analysis (PCoA) was performed on the rlog transformed sequencing data to visualize the broad trends of similarities and differences among samples ^54^. The first two coordinates and the explained variance were plotted on the two axes. To compare bacterial communities, we constructed a Bray-Curtis dissimilarity matrix based on the regularized logarithm (rlog) transformation counts of OTUs. Null hypothesis tests of no differences among *a priori*-defined groups were performed by permutational multivariate analysis of variance (PERMANOVA) ^47^. The significances of the treatments “days after rain” (fixed continuous covariate) and “land use” (fixed factor with two levels: urban vs residential) and their interaction were tested, accounting for flumes (random factor, six levels) (*alpha* = 0.05). Differences between “days after rain” and each “land use” were tested by pairwise comparisons, where significant interactions were observed. Relationships between changes in the microbial community structure and individual flume ecosystem descriptors were analysed using distance-based multivariate multiple regression (DistLM) ^47^. Two-sided significance tests determined whether a correlation was significantly different from zero (*p*-value < 0.05). Ecosystem descriptors were then subjected to a forward-selection procedure to develop a model to explain the variance in microbial community structure, taking into account “land use”, “days after rain”, and their interaction. Model selection was based on the Akaike information criterion, penalized for extra parameters (AICc) ^55, 56^.

## SUPPLEMENTARY INFORMATION

### SI Figures

**Figure S1.**
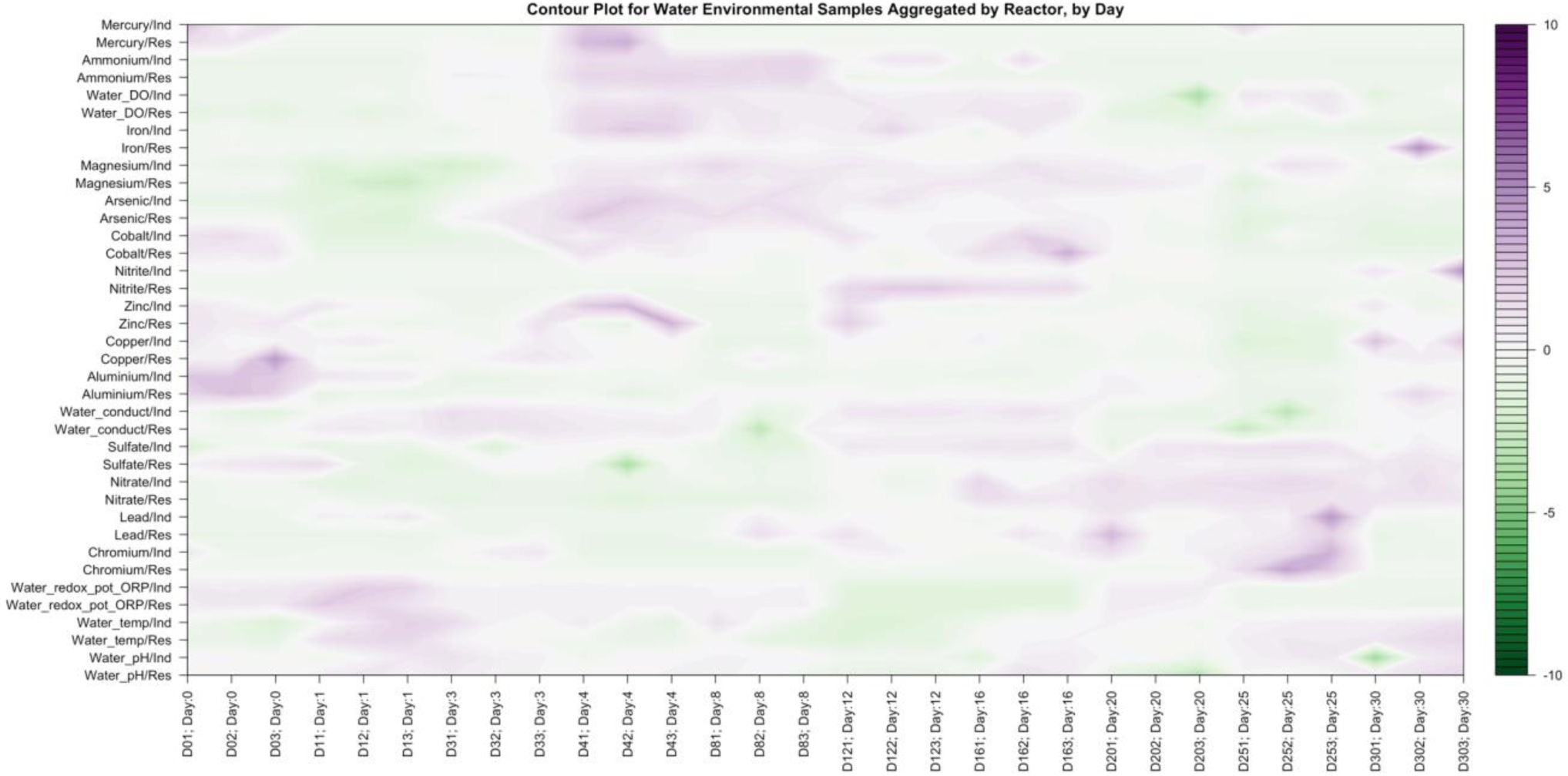
Contour plot of the environmental parameters relative levels in water samples collected during the 10 time-points spread over 30 days experiment.

**Figure S2.**
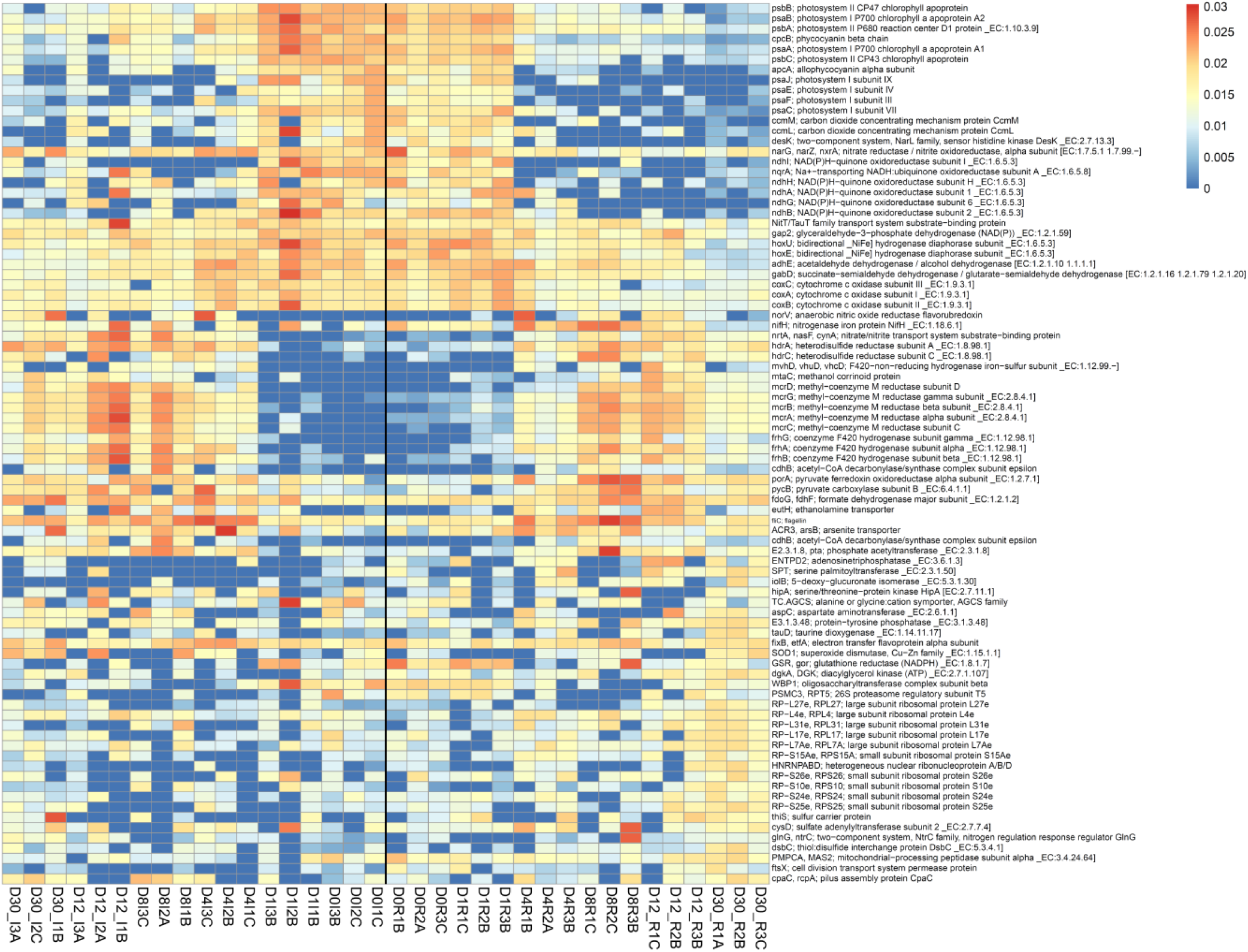
Heat map of the functional genes selected based on the high coefficient of variation.

### SI Tables

**Table S1:** PERMANOVA results on microbial community data derived from 16S amplicon sequencing, using Bray-Curtis distance similarity on regularized logarithm transformed (rlog) abundance data. Time and sediment type (land-use) were the fixed and orthogonal factors. Reactor number was random factor, nested in sediment type (land-use). ST: sediment type (land-use); T: Time (Days after rain); Re: Reactor; df: degree of freedom; SS: sum of squares; MS: mean sum of squares; Pseudo-F = F value by permutation

**PERMANOVA (taxa)**
|  | df | SS | MS | Pseudo-*F* | *p*(Monte-Carlo) |
| --- | --- | --- | --- | --- | --- |
| Time: Days after rain (T) | 9 | 36.55 | 4.06 | 7.53 | <0.001 |
| Sediment type: land-use (ST) | 1 | 5.56 | 5.56 | 10.29 | <0.001 |
| Flume (Re; nested in ST) | 4 | 2.16 | 0.54 | 1.00 | 0.481 |
| T * ST | 9 | 18.44 | 2.05 | 3.80 | <0.001 |
| Residuals | 36 | 19.43 | 0.54 |  |  |

Pairwise tests: All days: Industrial ≠ Residential; All sediment type (land-use): Days are
significantly different in the early and mid-phase of experiment.

Table S2 is available here: https://mega.nz/folder/zV02zKaT#70o6dxHwLkNa3erP8-zj5g/file/eYUVmaIJ

